# Structure adaptation in Omicron SARS-CoV-2/hACE2: Biophysical origins of evolutionary driving forces

**DOI:** 10.1101/2022.12.20.521221

**Authors:** Ya-Wen Hsiao, Tseden Taddese, Guadalupe Jiménez-Serratos, David J. Bray, Jason Crain

## Abstract

Since its emergence, the Covid19 pandemic has been sustained by a series of transmission waves initiated by new variants of the SARS-CoV-2 virus. Some of these arise with higher transmissivity and/or increased disease severity. Here we use molecular dynamics simulations to examine the modulation of the fundamental interactions between the receptor binding domain (RBD) of the spike glycoprotein and the host cell receptor (human angiotensin-converting enzyme 2: hACE2) arising from Omicron variant mutations (BA.1 and BA.2) relative to the original wild type strain. We find significant structural differences in the complexes which overall bring the spike protein and its receptor into closer proximity. These are consistent with and attributed to the higher positive charge on the RBD conferred by BA.1 and BA.2 mutations relative to the wild type. However, further differences between sub-variants BA.1 and BA.2 (which have equivalent RBD charges) are also evident: Mutations affect interdomain interactions between the up-chain and its clockwise neighbor chain, resulting in enhanced flexibility for BA.2. Consequently, additional close contacts arise in BA.2 which include binding to hACE2 by a second spike protein monomer, in addition to the up-chain - a motif not found in BA.1. Finally, the mechanism by which the glycans stabilize the up state of the Spike protein differs for the wild type and the Omicrons. We also found the glycan on N90 of hACE2 turns from inhibiting, to facilitating the binding to Omicron spike protein. These structural and electrostatic differences offer further insight into the mechanisms by which viral mutations modulate host cell binding and provide a biophysical basis for evolutionary driving forces.

## I. INTRODUCTION

Coronavirus disease-2019 (COVID-19) has been responsible for more than 6 million deaths and over 500 million cases worldwide over the course of the ongoing pandemic and it remains a significant global health threat. It is caused by the severe acute respiratory syndrome-coronavirus 2 (SARS-CoV-2) - a single-stranded RNA-enveloped virus and the seventh known coronavirus to infect humans. Since its wild type strain (WT) was first discovered in Wuhan, China, the virus has undergone continuous mutation leading to a succession of so-called variants of concern (VOC). Designated by the World Health Organization as Alpha to Delta, and Omicron, these VOCs occur with increasing transmission rate and variable pathogenicity.

In common with other coronaviruses, the virion surface consists of spike (S) glycoproteins comprising two (S1 and S2) subunits. The S2 subunit embeds the spike in the viral envelope and mediates viral fusion with the host cell membrane upon protease activation. The S1 subunit is involved in recognition and binding to the peptidase domain of the host receptor angiotensin-converting enzyme 2 (hACE2). It includes a trimer of N-terminal domains (NTD), and receptor binding domains (RBD) each of which exists in either of two discrete up or down conformations in the pre-fusion state. Only in the up-conformation is the receptor binding site exposed. Typically, in the active conformation one of the three RBD domains is in the up-state with the other two in the down position [1].

The, now dominant, Omicron strain is unique among known variants as it exhibits a much larger number of point mutations than did any of its predecessors - more than 30 on the S-protein, of which over half are found in the RBD. Since vaccine strategies rely heavily on the spike sequence, the emergence of such hypermutated strains threaten to weaken the neutralizing activity of vaccine-induced antibodies. Indeed, there is evidence that Omicron subvariants BA.1, BA.2 exhibit immune escape among double vaccinated hosts, and require a third dose to induce the neutralizing immune response [2, 3].

The molecular mechanisms by which the SARS-CoV-2 RBD recognizes and associates with hACE2 are therefore fundamental to the infection process. They are also essential for elucidating the biophysical consequences of mutation and, thereby, the molecular basis for evolutionary driving forces. For example, relative to WT, the net positive charge on each RBD increases by two and three for Delta and Omicron variants, respectively. This increase in charge is accompanied both by higher infection rate and reduced potency of neutralizing antibodies [4–12]. Both effects are consistent with evolutionary pressure toward RBD mutations which increase positive charge: Firstly, as hACE2 is a charge negative entity [13] thus the electrostatic interaction is increased; and secondly, the electrostatic surfaces of neutralizing antibodies reveal that most antibodies have positively charged RBD-recognition domains. However, so far reports generally have focused on binding free energy difference. Most of them were evaluated based on the binding structure of RBD+hACE2 for WT. Structural changes in the RBD+hACE2 complex arising from Omicron-lineage mutations have been less explored in comparison.

The effects of the individual point mutations on RBD with respect to infectivity have also been another focus of recent studies. For example, N501Y, the RBD mutation in the Alpha variant enhanced the binding affinity to hACE2 [14, 15]; E484K in Beta and Gamma variants can reduce the neutralization activity of human sera [16]; L452R in Delta and BA.2 variants have been shown to strengthen fusogenicity and infectivity by enhancing the cleavage of spike protein; and K417N removes the interfacial salt bridge between the RBD and neutralizing antibody CB6 [11]. Questions naturally arise concerning whether there are more ways to associate the high infectivity and the many point mutations on each RBD for BA.1/BA.2.

Fifteen and sixteen point mutations occur on each RBD of BA.1 and BA.2, respectively, and affect the interaction between RBD and hACE2. The mutations in the RBD of BA.1 and BA.2 contribute to an increase of the net positive charge by three compared to WT. A stronger electrostatic attraction of the RBD+hACE2 complex is thus expected for Omicron than WT. However, this leaves an obvious question as to why BA.2 is more infectious than BA.1 [17] as both have the same net charge at the RBD. Thus specifics of the localized electrostatic landscape must matter (i.e. interactions involving specific mutations) in binding the complex system.

Another key feature of the S-protein is its extensive glycosylation. Each monomer S-protein displays 22 potential N-linked and four O-linked glycosylation sites [18, 19]. Cryo-electron microscopy provides evidence for the existence of 17 N-glycans on 22 potential sites in the SARS-CoV-2 S-protein [1, 20], while the remaining five sites are found unoccupied [19]. These glycans shield about 40% of the trimeric S-protein surface [21], as well as other epitopes from cells and antibody recognition and enable the coronavirus to evade both the innate and adaptive immune responses [1, 22] Thus, glycosylation plays a vital role in the pathogenesis of virus such as coronavirus [18, 23, 24] Additionally, receptor hACE2 is also heavily glycosylated, and the glycan on N90 of hACE2 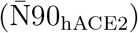 was found to be able to interfere with the binding between hACE2 and S-protein [25, 26], whereas 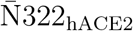 interacts tightly with the RBD of the hACE2-bound S-protein and strengthens the complex [26]. Therefore, it is evident that glycans, either on the S-protein or on hACE2, are an important factor for the formation of S+hACE2 complex.

Herein, we use molecular dynamics (MD) simulations to investigate the structural and biophysical mechanisms by which mutations in SARS-CoV-2 modulate receptor recognition and binding. We focus on the Omicron variant BA.1, and its sublineage BA.2 in complex with hACE2 and compared these to that of the WT. In addition to elucidating the effects of mutations in the binding of this complex system, we also examine the role of the glycans at the interface in forming S+hACE2 complex.

## II. METHODS

### A. Structural Model for WT, BA.1, BA.2 and Initial Configuration with hACE2

The structures for hACE2 and WT S-protein in the up state were modeled using the PDB structures 6VW1 [27] and 6VSB [20], respectively. The spatial arrangement of the three chains in the S-protein is shown in Figure 1. The color code of each S-protein chain is followed throughout this paper: chain A is the up-chain (cyan); chain B is the counterclockwise chain (yellow); chain C is the clockwise neighbor chain (lime). We obtained these from the CHARMM-GUI [28] Archive–COVID-19 Protein Library which arranged the position of hACE2 and S-protein, as given by Woo *et al*. [29]. Here, Woo *et al*. [29] modified the amino acid sequence of 6VSB to match the WT and had the remaining missing atoms of S-protein reconstructed.

**FIG. 1:**
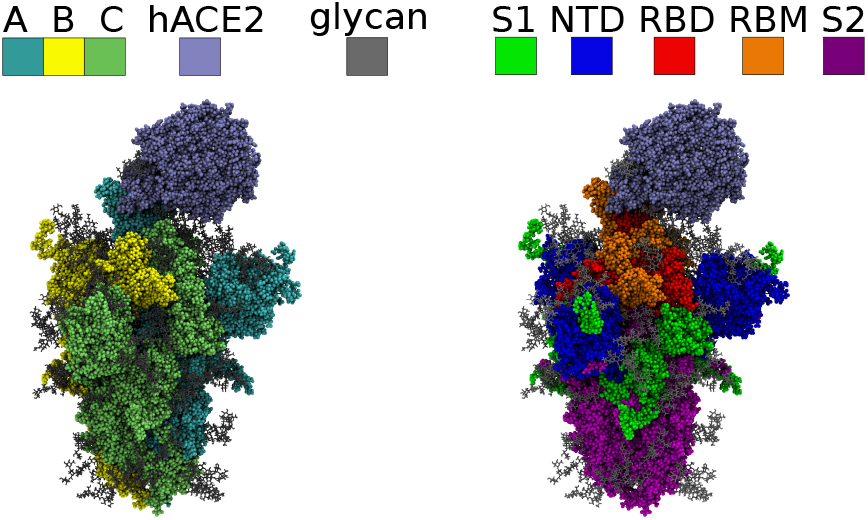
Representations of the quaternary structure of the spike protein and hACE2 highlighting the S trimer chains (A-C), key domain structure (RBD, RBM and NTD) and bound glycans.

Using the sequence of the WT S-protein as the template, we generated the Omicron subvariants BA.1 [30] and BA.2 [31] using VMD [32, 33] by introducing the corresponding sequence changes (using the VMD plugin psfgen). The full glycosylation model (for hACE2 and S-protein proposed by Shajahan *et al*. [34] and Woo *et al*. [29], respectively) was introduced to the protein structures using the Glycan Modeler within CHARMM-GUI[35].

Residues 19 to 614 of a monomer hACE2, and residues 1 to 1146 of trimeric S-protein, both fully glycosylated, were then solvated in a cubic box of TIP3P water [36] with 0.15 M NaCl and charge neutralized with additional Na ions (i.e., H_2_O/Na/Cl of 495442/1448/1406, 495060/1426/1405 and 495204/1426/1405, respectively, for WT, BA.1 and BA.2). Hence, resulting in initial box size of 255 Å *×* 255 Å *×* 255 Å (where the combined S + hACE2 structure takes up at most 234 Å *×* 170 Å *×* 158 Å).

### B. MD Simulation Protocal

All simulations were carried out using software NAMD 2.12 [37] and the CHARMM36 force field [38, 39]. First the initial system was energy minimized using steepest descent to remove atom overlap and then the solvent relaxed by running a MD simulation where the backbone atoms were restrained with a force constant of 5 kcal mol^−1^Å^−2^ for a simulated 10 ns duration. Afterwards, an unrestrained MD production run was performed for just over 1 μs. For each MD run, a 2 fs timestep was used with periodic boundary conditions and NPT ensemble (at 310 K and 1.01325 bar conditions) applied using a Langevin thermostat (1 ps^−1^ damping) and Nosé-Hoover Langevin piston barostat (50 fs period and 25 fs decay). The non-bonded interaction cut-off was set at 12 Å and switch distance at 10 Å. Bonds involving Hydrogen were made rigid using the SHAKE algorithm [40]. Electrostatics were calculated using Particle Mesh Ewald approach [41] (1 Å grid-spacing and 6 PME interpolation order).

To avoid translation, rotation and to limit swaying motions, part of the S2 subunit (over residues 701–729, 787–824, 866–939, and 1035–1146 on all three chains) was anchored via restraints on the backbone atoms using 20 kcal mol^−1^Å^−2^ force constant. Details in analyses the key measurements can be found in the supplemental information.

## III. RESULTS AND DISCUSSION

### A. Two types of S+hACE2 conformations

We found a trend change in the complex structures around 600 ns (see the left panel of Figure S2), thus, we analyzed the data from the last 400 ns of the simulation. The representative structures after 600 ns are displayed in Figure S1. Our results show two conformations for the S+hACE2 binding: one, an interface solely via RBD-A…hACE2 contact which we see for WT (i.e. contact with the up chain of S-protein); two, interfacing that involves both RBD-A…hACE2 and RBD-C…hACE2 contacts which we see for omicron variants BA.1 and BA.2 (i.e. contact involving the up chain and a down chain of S). We do not observe contact involving RBD-B.

#### 1. The Omicrons have smaller opening angles for the up state chain but larger opening angles for the down state chain C

We compared the up state structures of the three (sub)variants by measuring the opening angle, *θ*_RBD-A_, of the S-protein (an example of *θ*_RBD-A_, as used by Fallon *et al*. [42]), is shown in Figure 2). We found that the time averaged value for *θ*_RBD-A_ ranked WT *>* BA.2 *>* BA.1 (Table I, left panel of Figure S2). We also measured the opening angle *θ*_RBD-C_ to check whether chain C becomes open in order to bind hACE2 (Table I, right panel of Figure S2). We find that the Omicrons have larger *θ*_RBD-C_ than WT (i.e., RBD-C is more open for Omicrons with BA.2 being greater than BA.1), but are much less open than RBD-A, where *θ*_RBD-A_ *>* 75° compared to *θ*_RBD-C_ *<* 60° (i.e., RBD-C is never observed in the fully open state). For completeness we measured *θ*_RBD-B_ and obtained average values for WT: 70.5°; BA.1: 52.6°; BA.2: 52.2°. This result implies that RBD-B of BA.1/BA.2 is also closed.

**TABLE I:** Average RBD opening angle (°); and radius of gyration R_*gyr*_ (Å)

| Variant | $\theta_{\text{RBD-A}}$ | $\theta_{\text{RBD-C}}$ | $R_{gyr}^S$ | $R_{gyr}^{S+hACE2}$ |
| --- | --- | --- | --- | --- |
| WT | $88.7 \pm 1.8$ | $45.2 \pm 1.1$ | $51.6 \pm 0.1$ | $60.8 \pm 0.3$ |
| BA.1 | $75.1 \pm 1.7$ | $55.8 \pm 1.4$ | $51.2 \pm 0.1$ | $60.0 \pm 0.3$ |
| BA.2 | $83.1 \pm 1.4$ | $58.9 \pm 1.8$ | $51.3 \pm 0.1$ | $59.3 \pm 0.2$ |

**FIG. 2:**
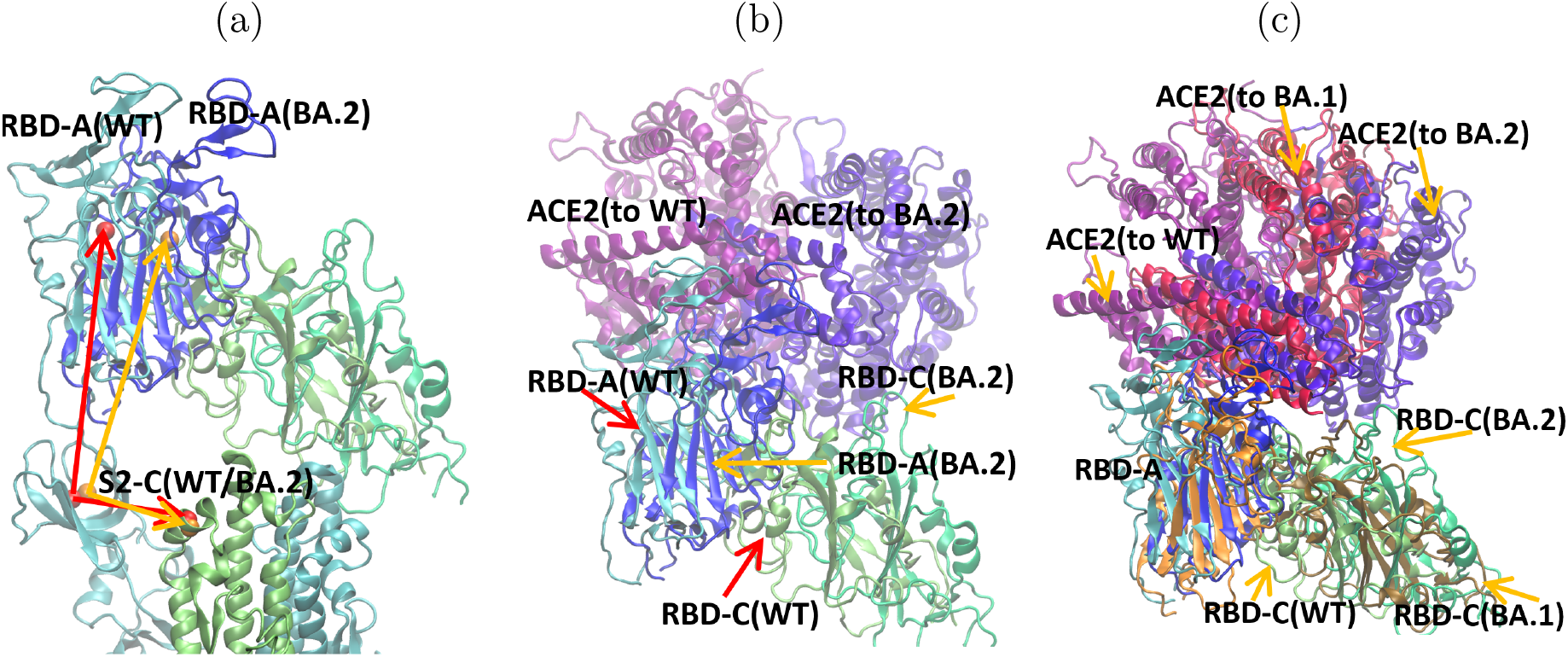
Superimposed images of WT, BA.1 and BA.2 aligned along the S2 sub-unit: (a) definition of *θ*_RBD-A_ for WT (red arrows) and BA.2 (orange arrows); (b) identification of RBD-A, RBD-C of WT and BA.2, with the connected hACE2; (c) BA.1 and the corresponding hACE2 are also included.

#### 2. The Omicron variants have more compact S-protein + hACE2 complexes

To understand why *θ*_RBD-A_ varies we measured the radius of gyration of the trimeric S-protein, 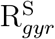, and the combined S + hACE2 structure, 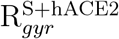 (Table I, Figure S3(a)). For the Omicrons, 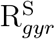 shows only a slight decrease, by 0.3-0.4 Å, compared to WT, whereas for 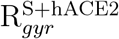 BA.1 decreases by 0.8 Å and BA.2 by 1.5 Å compared to WT. The smaller 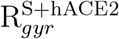 for BA.2 implies a more compact structure and suggests that BA.2 is the most tightly bound to hACE2 of the three variants.

We investigated the center of mass (COM) distances 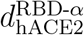 to find out how each individual chain *α* contributes to the change in 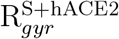 (see Table II, Figure S3(b-d)). Clearly, BA.2 has the shortest distance between hACE2 and any of its three RBDs. For RBD-A, BA.1 is the longest, but not able to explain the slightly shorter R_*gyr*_ for BA.1 compared to WT without taking into account the contribution of the other two RBDs. 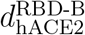 and 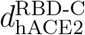 share the same trend: WT *>* BA.1 *>* BA.2, same as seen in 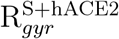. The differ-ence is distinctly larger for RBD-C: 4.8 Å (BA.1) and 10 Å (BA.2) compared to WT. This substantial shortening of 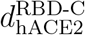 can increase the chance of RBD-C to interact with hACE2. Further, the above mentioned smaller *θ*_RBD-A_ of BA.1/BA.2 is due to the change of relative position. The middle panel of Figure 2 shows that hACE2 changes its orientation from binding to WT to BA.2 (S2 of WT and BA.2 S-proteins aligned). This is the origin of the smaller *θ*_RBD-A_ as well as the closeness of hACE2 to RBD-C. The right panel of Figure 2 includes also the RBD-A, RBD-C, and corresponding hACE2 of BA.1, allowing one to see the gradual change of the relative position of hACE2 from WT, BA.1, to BA.2.

**TABLE II:** COM distance (Å) between each chain RBD domain and hACE2

| Variant | $d_{hACE2}^{\text{RBD-A}}$ | $d_{hACE2}^{\text{RBD-B}}$ | $d_{hACE2}^{\text{RBD-C}}$ |
| --- | --- | --- | --- |
| WT | $48.8 \pm 0.5$ | $77.6 \pm 2.3$ | $61.0 \pm 1.8$ |
| BA.1 | $50.3 \pm 0.7$ | $76.3 \pm 1.5$ | $56.2 \pm 1.6$ |
| BA.2 | $46.9 \pm 0.5$ | $72.7 \pm 1.0$ | $51.0 \pm 0.4$ |

To understand what drives the S+hACE2 complex into the two observed conformations described we performed a detailed study into the contact interactions. Contact comes in two forms: direct interaction between amino-acids and indirect interaction through the selected glycans attached to S-protein and hACE2. In the next few sections we will look at each in turn.

### B. Direct interaction between S-protein and hACE2

#### 1. Variation in key contacts hACE2…RBD-A involving mutated residues

From our simulation trajectories we identified the contacts between RBD-A and hACE2. One might assume that the positively charged mutations on the RBD (WT → Omicron) enhance the RBD-A binding with hACE2. However, our results showed it more likely, that the mutations change the contact partners on hACE2, as summarized in Table III. For BA.1, we find that the our contact pairs are consistent with those found by Lan *et al*. [43], e.g., the interactions between BA.1 RBD residues (Q498R and N501Y) and hACE2 residues Y41 and Q42, as well as the loss of interactions between K417N(BA.1) and D30(hACE2); Y505H(BA.1) and R393/E37(hACE2); Q493R(BA.1) and K31(hACE2). After mutation, new contacts with hACE2 are made involving G496S_A_ for BA.1, N440K_A_ and D405N_A_ for BA.2. Although N440K_A_ is common to BA.1 and BA.2, it interacts with E329_hACE2_ more strongly for BA.2 (90%) than BA.1 (27%); instead N440K of BA.1 is fully solvated by water for the most of the time.

Table III also shows that the RBD-A…hACE2 contacts are rather dynamic, since there can be more than one residue with high probability of making close contact. Experimental assays and protein structure analysis by Bhattacharjee *et al*. [44] showed that hACE2 interfacial residues can bind to multiple S-protein residues, for example, K353_hACE2_ can interact with as many as six S-protein residues Y505, N501, G496, Q498, G502, and Y495. We see K353_hACE2_ interact with G496/G496S, N501/N501Y, and Y505/Y505H, as well as Q493R (but not Q493 WT).

**TABLE III:** Most probable contact pairing between key mutated residues of RBD-A and hACE2 with % occurrence given in brackets. Entries in bold indicate contact involving the mutated residue.

| RBD-A Residue | WT | BA.1 | BA.2 |
| --- | --- | --- | --- |
| K417N | D30 (90%) | – | – |
| N440K | – | **E329 (27%)** | **E329 (90%)** |
| S477N | T20 (7%) | **Q24 (3%)** | **S19 (13%)/T20 (10%)** |
| T478K | Q24 (7%) | **Y83/P84 (2%)** | **Q24 (9%)/P84 (6%)** |
| E484A | K31 (17%) | **Q76/L79 (37%)** | **L79 (87%)** |
| Q493R | E35 (77%)/K31 (60%) | **D38/H34 (100%)/K353 (95%)** | **D38/H34 (99%)/K353 (77%)** |
| G496S | K353 (23%) | **K353 (77%)** | – |
| Q498R | Y41 (81%)/Q42 (41%) | **Y41 (94%)/Q42 (57%)** | **Y41 (90%)/Q42 (37%)** |
| N501Y | K353/D355 (100%)/Y41 (86%) | **K353/D355 (100%)/Y41 (77%)** | **K353/D355 (100%)/Y41 (30%)** |
| Y505H | K353/R393 (99%)/E37 (100%) | **G354/K353 (98%)** | **A386/A387 (98%)/K353 (63%)** |
| D405N | A387 (3%) | – | **A387 (71%)** |
| R408S | A387 (3%) | – | – |

The mutations D405N and R408S are unique to BA.2 as they are not present in BA.1. D405N eliminates one negative charge and R408S eliminates one positive charge, therefore the net charge on the RBDs is the same between BA.1 and BA.2. However, while D405N_A_ contributes substantially to the interaction with hACE2 (Table III, Figure 3b), R408S_A_ does not participate in hACE2 binding. This is consistent with the finding of Li *et al*. [45] who obtained similar binding affinities for mutants with R408S and S408R and concluded that the R408S mutation contributes very limited RBD binding to hACE2.

**FIG. 3:**
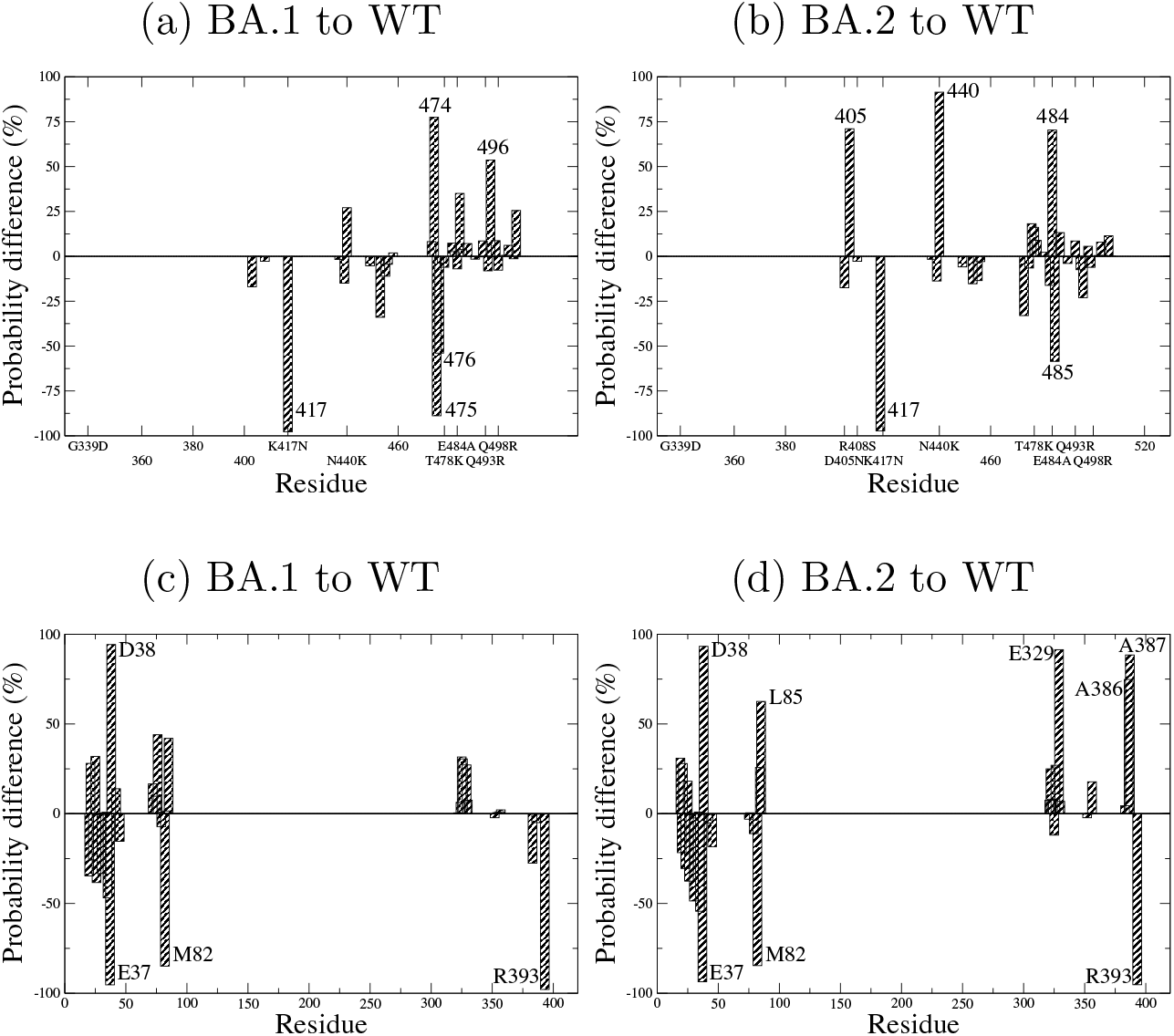
Probability difference of RBD-A residues of S-protein in contact with hACE2 ((a), (b)); and hACE2 residues in contact with RBD-A of S-protein ((c), (d)). Residues contributing more than 50% are labeled.

The contact probabilities are plotted in Figure S4, and additional contact details are collected in Table S1 including contact pairs for residues that are not mutated.

#### 2. The difference in contact probability of RBD-A…hACE2 between variants

With respect to the residues on RBD-A, we evaluated the difference in the probability of contact between RBD-A and hACE2 for the two Omicron subvariants against WT, as depicted in Figures 3a, 3b.

We find that contacts involving the above mentioned RBD-A residues G496S_A_ for BA.1 and D405N_A_, N440K_A_ for BA.2 show a large increase in occurrence after mutation, (54%, 71%, 91% respectively). For both BA.1 and BA.2, the mutation at K417N_A_ results in the loss of contact to hACE2 (a 98% reduction in contact with D30_hACE2_). This observation is consistent with Luan and Huynh [11] who suggest that this may be for eluding antibodies. For BA.1 (Figure 3a), A475_A_ and G476_A_ also lose contact with hACE2 substantially (89% and 54%, respectively), but is replaced by a new nearby contact at Q474_A_ (to Q24_hACE2_). In BA.2 E484A_A_, another common mutation for BA.1 and BA.2, takes over the contact to L79_hACE2_ from G485_A_ of WT and BA.1, thus showing another example of changing contact partner.

To determine whether the S-protein mutations affect the accessibility of RBD-A to hACE2, we evaluated the probability difference of residues on hACE2 in close contact with RBD-A (Figures 3c,3d). Compared to WT, the contact probability for hACE2 to RBD-A is slightly lower for BA.1 (overall probability ratio 0.94) but slightly higher for BA.2 (overall probability ratio 1.05). The residues of contact on hACE2 differ between BA.1 and BA.2 mainly at E329_hACE2_ and A386/A387_hACE2_ where BA.2 establishes close contact using N440K_A_ and D405N/Y505H_A_.

There is a probability decrease at E37_hACE2_ and increase at D38_hACE2_ which is caused by the contact between E37_hACE2_ and Y505_A_ of WT seemingly being eliminated in BA.1/BA.2 by the mutation Y505H_A_, and by a salt bridge Q493R_A_…D38_hACE2_ forming in preference to E37_hACE2_. This change of binding pattern was also found by Lupala *et al*. [46], in the case of mutation Q493K (rather than Q493R seen here). Besides binding to E37_hACE2_, Y505_A_ of WT additionally binds to R393_hACE2_, whereas for BA.1/BA.2 the mutated Y505H_A_ leaves R393_hACE2_ largely solvated by water and rarely connected to hACE2. Furthermore, M82_hACE2_ becomes water-exposed in BA.1/BA.2, and the contact L85_hACE2_…F486_RBD-A_ is 42% and 63% more likely to occur for BA.1 and BA.2, respectively, than for WT.

The interactions between E329/A386/A387_hACE2_ and RBD-A are unique for BA.2, and may be relevant to the difference between BA.1 and BA.2: the fact that more interfacial hACE2 residues are in close contact to RBD-A of BA.2 also suggests that BA.2 is more tightly bound than the others, consistent with its smaller 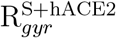. We calculated the average distance between the lysine sidechain nitrogen atom of N440K_A_ and the sidechain carboxylic carbon atom of E329_hACE2_ as 5.8 Å and 3.6 Å for BA.1 and BA.2, respectively.

The longer distance for BA.1 is consistent with the hypothesis that RBD-A of BA.2 is more tightly bound to hACE2 than that of BA.1.

#### 3. RBD-C of BA.2 forms contacts to hACE2

Based on the much shorter 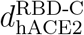 for BA.2, we hypothesized that its RBD-C can interact with hACE2, and carried out a similar probability analysis (as done for RBD-A) to see whether any contacts between RBD-C and hACE2 exists. We found that there is practically no close contact from WT (probability of contact totalling *<* 0.05%), and slightly more for BA.1 (*<* 1.5%), while substantial amount of interaction is found for BA.2 (*>* 99% for residues 445-449 and 498; 30–40% for residues 483-484). The results are shown in the upper row of Figure 4 where the major binding partners on hACE2 are residues 212-215, 564, 568, and 571-572 (lower row of Figures 4). These residues are different from those that bind to RBD-A (Table S1, Figures 3c, 3d), and therefore contribute additional binding between hACE2 and S-protein.

**FIG. 4:**
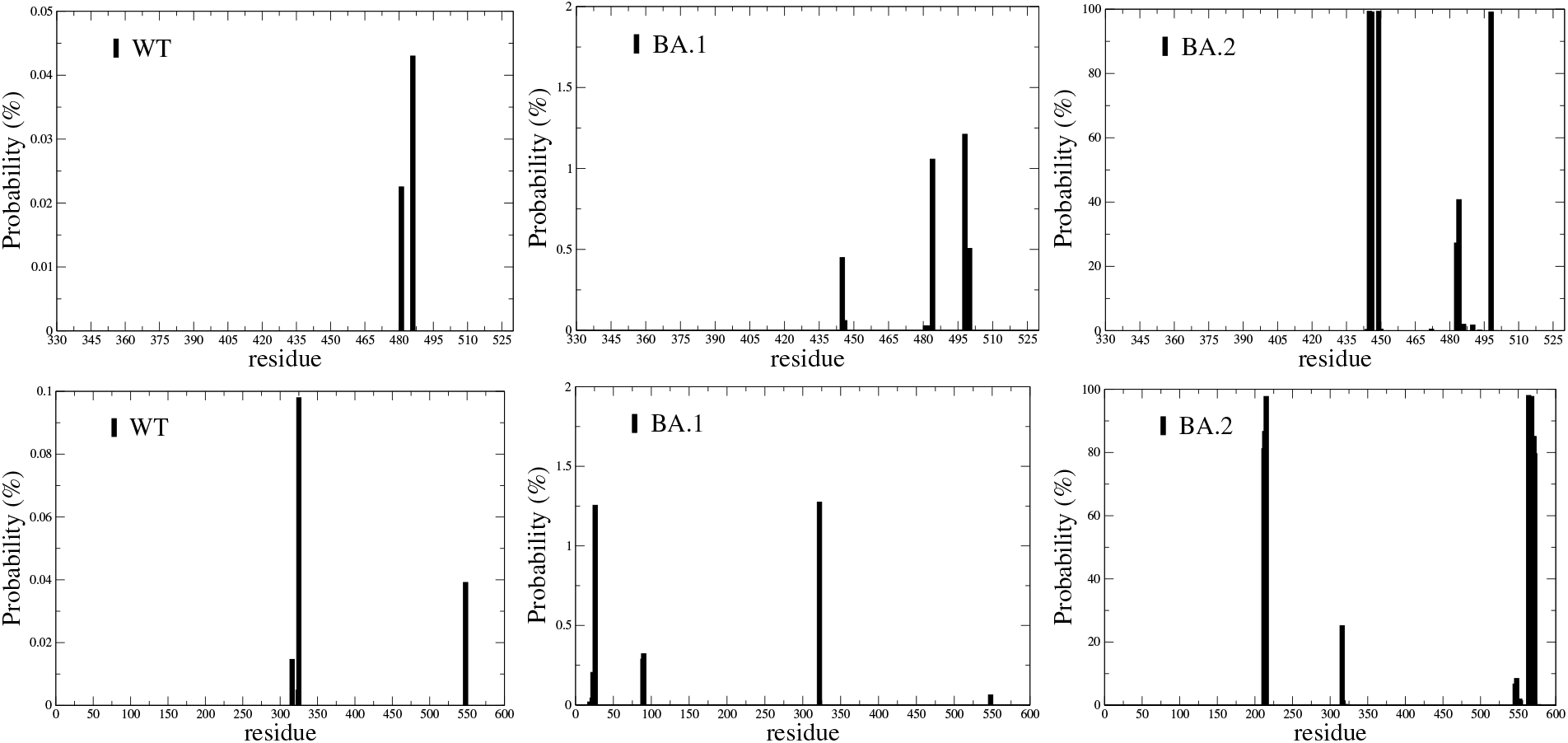
Probability of RBD-C residues in contact with hACE2 (top) and probability of hACE2 residues in contact with RBD-C (bottom). Note the different y-axis scales with BA.2 being much larger than either BA.1 or WT.

Multiple-RBD binding on one monomeric hACE2 for the Omicrons was qualitatively inferred in Yin *et al*. [47] by the observation that a monomeric hACE2 binds to a trimeric Omicron S-protein with 6-fold higher binding affinity than to WT S-protein, and only 2-fold higher affinity when binding to an Omicron monomer S-protein than to WT. As we did not model monomeric hACE2 interacting with a monomer S-protein we cannot make the same comparison here but nevertheless such scenario is consistent with our results.

### C. Degree of Interaction between RBDs

#### 1. Increased flexibility of RBD-C in BA.2

From the above, arises the question: why can RBD-C form contacts with hACE2 for BA.2 but not for BA.1 or WT? In Figure 5 we show the RMSF of the RBD residues for each chain of S. For RBD-A, we see that the RMSF of BA.2 is slightly higher (more flexible) of the three variants (except over the loop consisting of residues 476 to 485 where the RMSF is highest for BA.1). For RBD-B, the RMSF of BA.1 is the lowest (the chain is the lest flexible) and WT’s value is generally larger (more flexible) than BA.2, except over residues 476 to 485. For RBD-C, we find that BA.2 has the highest RMSF (showing that it is the most flexible) and WT the least.

**FIG. 5:**
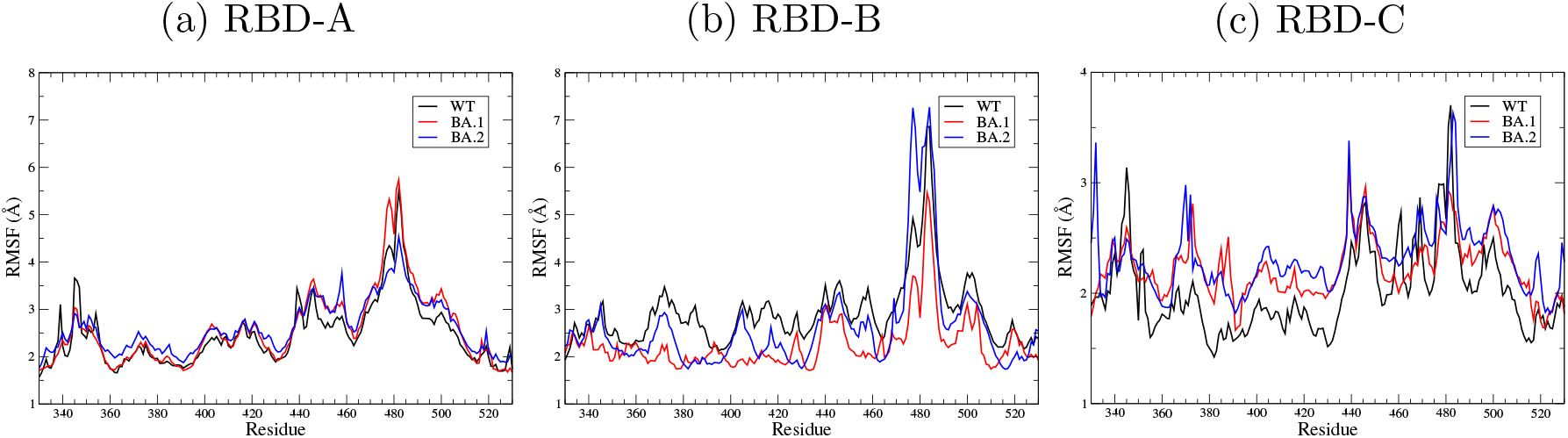
RMSF profiles for RBDs of S-protein.

#### 2. RBDs of chain A and C are more decoupled for BA.2

From the RMSF, we hypothesized that RBD-C’s ability to form contacts with h-ACE2 has to do with the increased freedom of RBD-C to restructure due to having less contact with the other RBDs. We hence investigated the degree of interaction between the RBDs and searched for key differences due to the mutations. Figure 6 shows the probability change in the inter-domain RBD-A…RBD-C contact from WT to BA.1/BA.2 (the full contact probabilities are shown in Figure S5). While the overall probability ratio of BA.1 over WT is around one, it is 0.44 for BA.2. Thus, there is a decoupling of the RBD-A and RBD-C in BA.2.

**FIG. 6:**
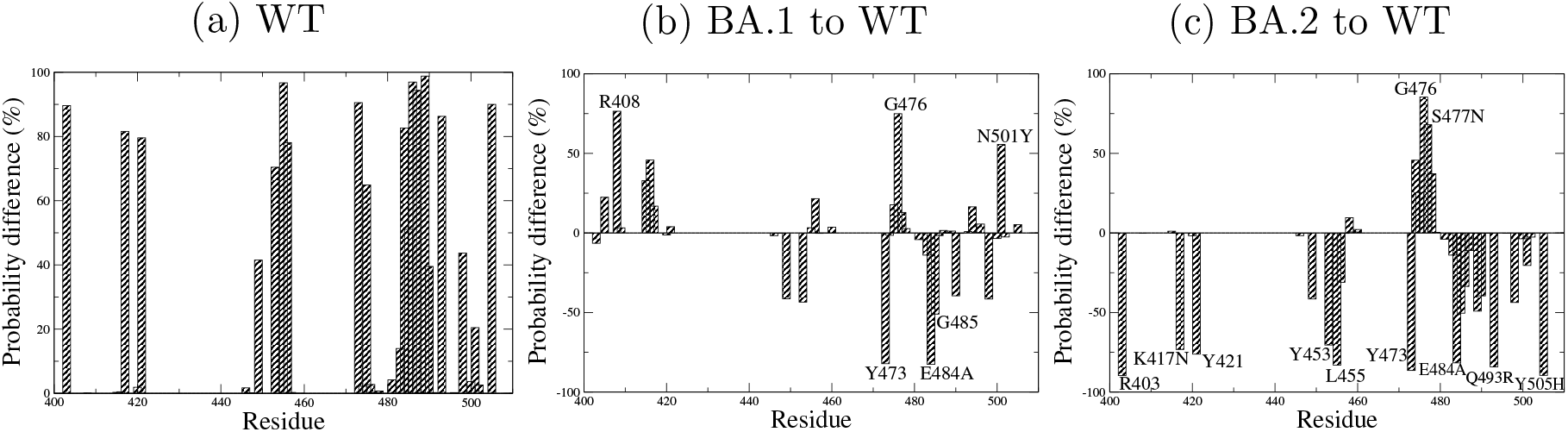
RBD-C residues in contact with RBD-A: probability of WT and probability difference of BA.1/BA.2 to WT. Residues differing by more than 50% are labeled.

This is due to several differences in RBD-A…RBD-C contact between the varients, over the regions 400-420_C_ and 440-500_C_. The latter is within the RBM_C_ and is also exposed to hACE2. For region 400-420_C_ there is an enhancement of interactions for BA.1 but loss of interactions for BA.2. In none of our models are there any contacts with hACE2 within this region. Here, BA.1 exhibits an additional contact made with R408_C_ (and to a lesser extent N501Y_C_) which is not present in either BA.2 or WT. Furthermore, for BA.1, R408_B_ forms contacts with residues 374 to 377 on RBD-C (a 98% presence). These contacts are missing in both BA.2, which has the mutation of R408S, and WT. We note that R408_C_ is located in the inner cavity formed by the three RBDs. In the case of BA.1, D428_A_ and R408_C_ form a salt bridge at the probability of 73%. This contact to D428_A_ is nearly absent for BA.2 because of the R408S mutation. For WT, due to the different relative position of RBD-C compared to the Omicrons, instead of R408_C_, D428_A_ connects to Y505_C_ (85%).

For region 440-500_C_, i.e., RBM_C_, there is an overall reduction in RBD-A…RBD-C interactions between Omicrons and WT which corresponds to the increase in interaction RBD-C…hACE2, as seen earlier, suggesting that RBD-C is pulled away from inter-chain S contact towards hACE2. By inspecting the solvent exposed side of RBD-A and RBD-C, we find that a salt bridge by K378_A_…E484_C_ is the dominant interaction (92%), and a hydrogen bonding pair S375_A_…E484_C_ (31%) forms at this interface for WT. Mutation E484A results in the loss of this salt bridge (shown in the probability difference in Figure 6) and gives further flexibility for RBD-C for both BA.1 and BA.2. E484A of BA.2 can in fact moderately approach hACE2 to residues V316 and T548 (29%), which is not seen for WT or BA.1. E484A loosens one connectivity between RBD-A and RBD-C for both BA.1 and BA.2, whereas the R408S mutation on RBD-C of BA.2 further loosens two connections of RBD-C: one to RBD-A and one to RBD-B, thereby setting BA.2 apart from BA.1, and gives BA.2 the most flexible RBD-C. The lower row of Figure 7 depicts the contact pair D428_A_…Y505_C_ in WT which is replaced with the new pair D428_A_…R408_C_ of BA.1 (with the mutation N501Y/Y505H as well), but this close link between RBD-A and RBD-C is loosened (due to R408S) for BA.2.

**FIG. 7:**
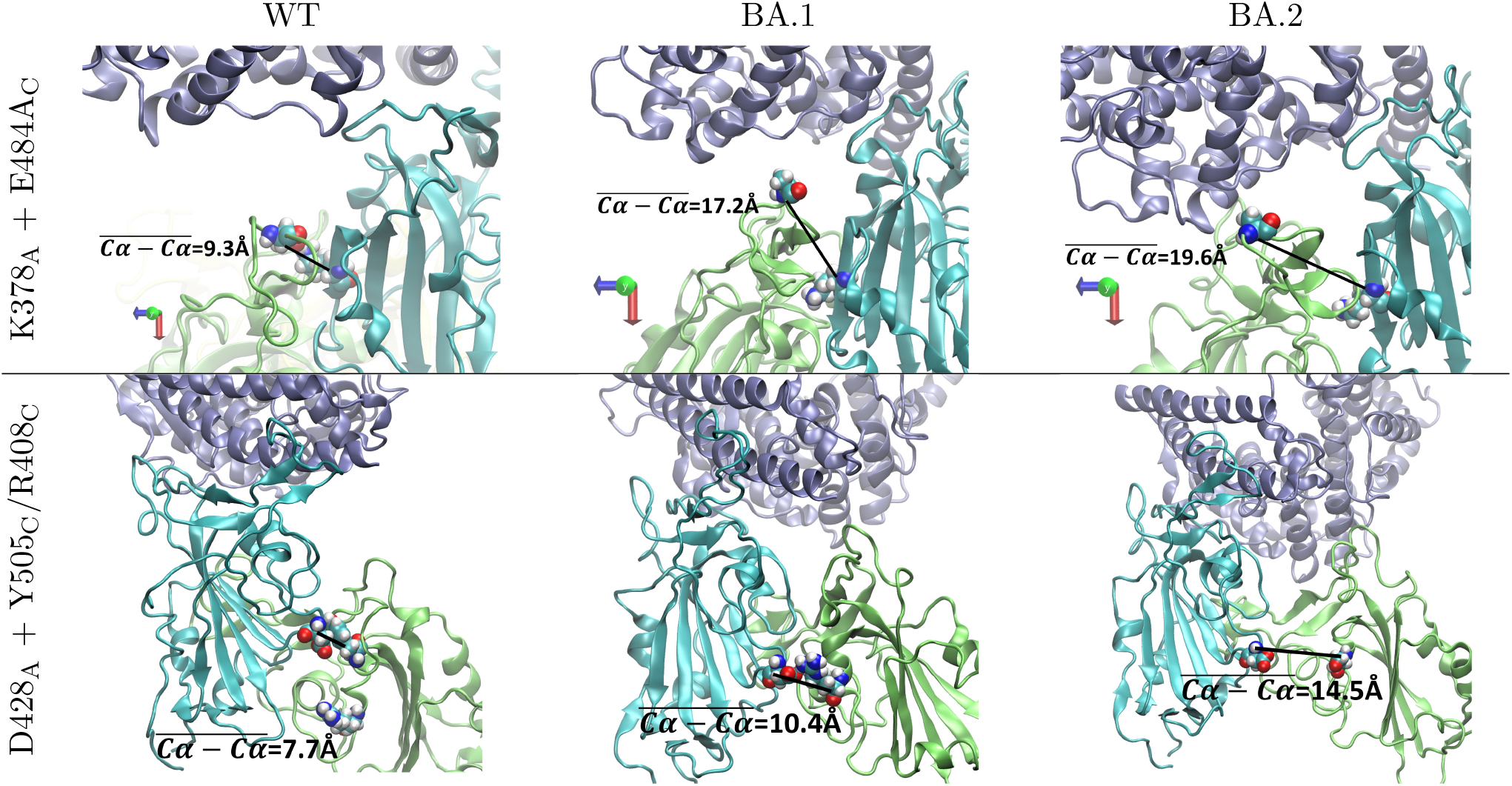
Relative position of key inter-domain residue pairing (‘ball-on-stick’) of S-protein’s RBDs (RBD-A - cyan, RBD-C - lime). hACE2 (lilac) is also shown. K378_A_…E484_C_ represents a salt bridge for WT, which is lost for the Omicrons because of the mutation E484A. D428_A_ contacts via the hydrogen bond D428_A_…Y505_C_ for WT, and salt bridge D428_A_…R408_C_ for BA.1 but is free for BA.2.

### D. The role of glycans in interfacing

The last part of the puzzle of how hACE2 interfaces with S-protein in our models lies with the role glycans play in the interaction. Of particular interest are 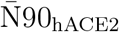 and 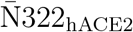 near the interface between hACE2 and S-protein. These two glycans of hACE2 are predicted to form interactions with S-protein [26, 34]. However, although 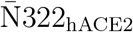 is in the vicinity of the RBD, its interaction with a monomeric S-protein was shown to be transient [48]. Moreover, Shajahan *et al*. [34] also pointed out that site N322 is aligned in human, bat, cat, pig and chicken, but pig and chicken are not susceptible to SARS-CoV-2 binding. In contrast, species that are susceptible have an N90 site while non-susceptible species do not [34], and 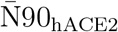 was shown to contact RBD with high probability [48].

Our analysis on the contact frequency between 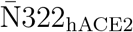 and RBD-A/RBD-C shows dominant interaction with RBD-C across all variants: 97%, 99%, and 92% for WT, BA.1, and BA.2, respectively. Whereas for RBD-A, contacts are formed at 13%, 0.4%, and 94% of the total occurrences for WT, BA.1, and BA.2, respectively. The only conclusion we can draw here is that 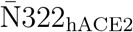 has stronger interaction with BA.2. The not so decisive role of 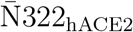 is probably due to its location which is outside the RBDs. On the other hand, 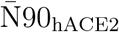 interfaces to RBDs directly. Here we will therefore observe the behaviors of 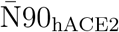, in our models of WT and the Omicrons and discuss its involvement in mediating the interactions between hACE2 and S-protein.

#### 1. 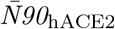 regulates/mediates the interfacing between hACE2 and S-protein

Previously, Mehdipour and Humme [26] have shown that 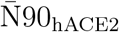 interferes with the binding between hACE2 and S-protein. While, Chan *et al*. [49] designed a decoy soluble hACE2 mutated at N90 to remove its N-glycan, and obtained high binding affinity to S-protein. To understand whether our models show 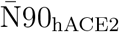 interfering with the binding of S+hACE2 we measured the orientation angle (Θ_N90_) of 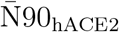 with respect to the interface (Figure S6). As can be seen from Figure 8 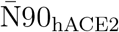 is most parallel to the S+hACE2 interface when binding to BA.2 (average Θ_N90_ = 7.5°), less so for BA.1 (Θ_N90_ = 35.6°), and more orthogonal for WT (Θ_N90_ = 100.5°).

**FIG. 8:**
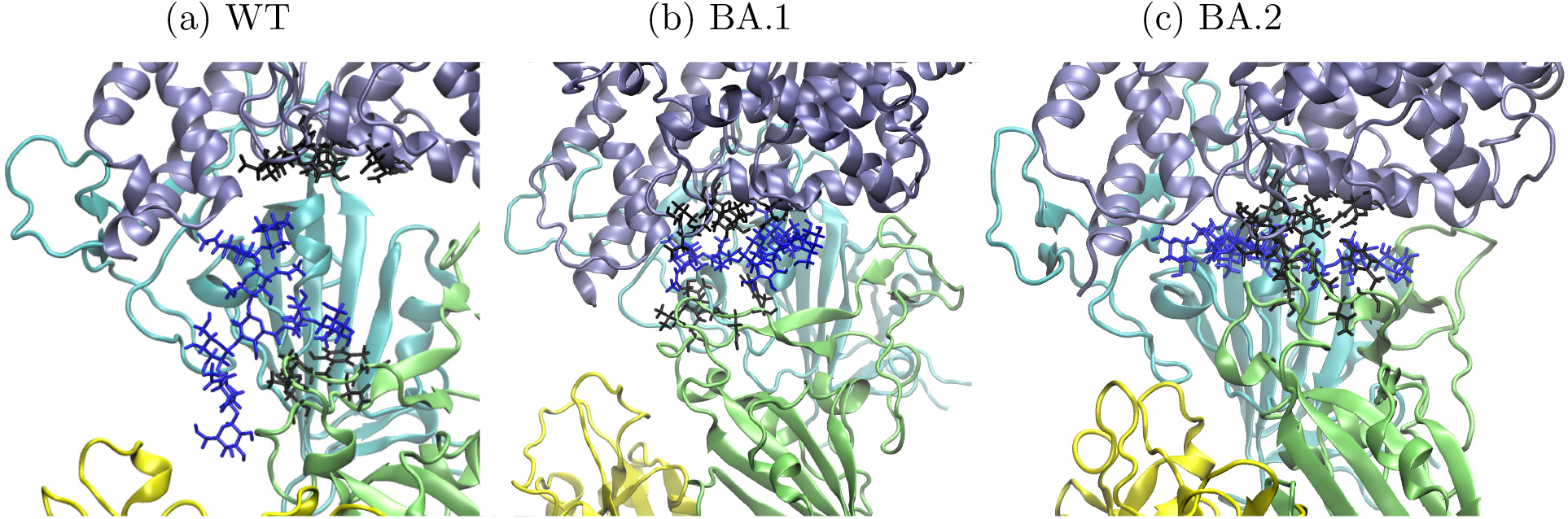
Relative position of 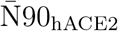 (blue licorice) to hACE2 (lilac) and RBDs (RBD-A - cyan, RBD-B - yellow, RBD-C - lime). The side-chains of RBD-C residues involved in binding with hACE2 are shown as black licorice.

Figure S7 shows the probabilities of RBD residues in contact with 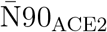. RBD-A’s of WT and BA.1 have little contact with 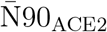, only N460 has a *>*50% contact presence for the former, and R408 the latter. N460 sits in the inner cavity, whereas R408 is half way to the solvent-exposed side, so that the main contact region on RBD-A differs between WT and BA.1. In contrast, substantial contacts to 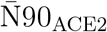 by RBD-A of BA.2 viz. D405N, R408S, Q409, T415, and G416, wherein D405N and R408S are away from the inner cavity. By having contacts spreading from inner cavity towards the solvent-exposed side, BA.2 keeps 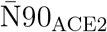 more parallel to the S–hACE2 interface thereby orienting 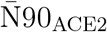 at the interface between RBD and hACE2.

Regarding RBD-C, for the Omicrons and particularly for BA.2, the contacts spread more extensively over RBM_C_ compared to WT. The residues forming good contact (*>*50%) with N 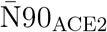 are: V445 and T500 for WT; V445, G446S, Y449, N450, and Q498R for BA.1; G447, Y449, L452, Q493R, S494, G496, Q498R, T500, N501Y for BA.2, wherein T500 and N501Y (inner cavity), as well as S494 (further out), maintain their contact throughout the simulation. These contacts of BA.2 to 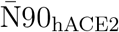 thus brings RBM_C_ to the vicinity of hACE2 as well as aligns RBM_C_ to the interface of S–hACE2 and eventually results in binding to hACE2 without the need of opening RBD_C_.

Owing to the few and inner-cavity-only contacts to both RBD-A and RBD-C, 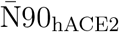 drops vertically (orthogonal to the S+hACE2 interface) for WT so that the space between hACE2 and S-protein is comparatively large, reflected in its large COM distance 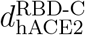 (Table II), and consistent with experimental observation of Mehdipour and Humme [26]. Whereas for BA.1/BA.2, more so for BA.2, this space is reduced which is consistent with the smaller 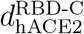. Here, the long axis of 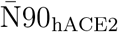 poses parallel to the S–hACE2 interface due to the contacts along the interface to both RBD-A and RBD-C. Therefore, for BA.2, 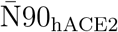 no longer acts as a spacer between RBD and hACE2. Compared to BA.2, BA.1 has fewer and less widespread contacts to, presumably due to the above-mentioned higher degree of RBD-A…RBD-C interaction resulting in lower RBD-A/RBD-C flexibility of BA.1.

## IV. CONCLUSION

In this work, we used MD simulations to investigate the SARS-CoV2 S+hACE2 structures, with S-protein from WT, and the Omicron subvariants BA.1, and BA.2. Starting from the open conformation given by Woo *et al*. [29], we find two conformations for the interfacing of S-protein and hACE2. A more open contact involving only predominately chain A (seen for WT and being of the same form as the starting structure) and a more compact conformation involving both chains A and C (seen for the Omicrons). The key feature of this interface is the injection of the glycan 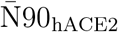 between hACE2 and S-protein and highlights the important interplay between direct binding of S-protein and hACE2 residues and indirect binding via interactions through 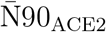 (i.e., 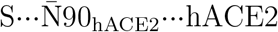). The key factors we found that promote this binding mode are as follows:

- Change in the S…hACE2 interface:
  − mutations in RBD increase the total positive charge which increases the electrostatic attraction between S-protein and hACE2. Nevertheless, we observed shifts in the binding partners after mutation.
  − BA.2 forms a broad contact with 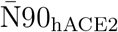, spreading from inner cavity towards the solvent-exposed side (Figure 8c), and thereby reducing the interfacial gap and holding hACE2 close to the S-protein to bind.
- One neighbor to the up-RBD, RBD-C, is more flexible for the Omicrons by less coupling to other RBDs, hence free to approach hACE2. This is more obvious for BA.2 than BA.1:
  − E484A of BA.1 and BA.2: abolishes a salt bridge K378_A_…E484_C_ at the solvent facing side in WT
  − R408S of BA.2: abolishes a salt bridge D428_A_…R408_C_ toward the inner cavity in BA.1, as well as cuts the connection with RBD-B (e.g., R408_B_…F374_C_ in BA.1), and therefore further reduces the connectivity to RBD-C.

The mutations E484A, R408S, and D405N in our result affect the structure, and the latter two seemingly make BA.2 distinct from others in the study. Although there is a loss of positive charge in R408S which is regained via the mutation D405N and keep the level of electrostatic interaction. These mutations are maintained in the later, and more infectious, subvariants BA.4 and BA.5, suggesting that these are indeed favorable mutations for increasing fitness of the virus. Moreover, the high degree of interaction between glycans on hACE2 and S-protein indicates that it is advantageous to consider such glycan-related epitopes in the therapeutic design. It has been suggested that the increased electrostatic attraction between S-protein and hACE2 may simply enhance the binding. Here we find that the binding can be further strengthened by the binding mode involving multi-RBDs per hACE2 monomer the Omicrons tend to adopt.

The structural reports introduced and investigated here can be more generally used to monitor emerging variants and future structural evolution of the virus and thereby developing a better model of the biophysical basis for evolutionary advantage.

## Supporting information

Key Measurements Made and Supplemental Table S1 and Figures S1-S7

final structure of S+hACE2 of the WT

final structure of S+hACE2 of BA.1

final structure of S+hACE2 of BA.2

## ACKNOWLEDGEMENTS

This work was supported by the Hartree National Centre for Digital Innovation, a collaboration between STFC and IBM.

## SUPPORTING INFORMATION

Key measure details, representative S+hACE2 complex structures, opening angle of RBD, radius of gyration, distance between center of mass, contact probabilities between RBD-A and hACE2, as well as between RBD-A and RBD-C, glycan orientation with respect to the hACE2-RBD interface, and contact probabilities between RBD-A/RBD-C and glycan 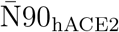 are collected in the supporting information (PDF), as well as the final structures of S+hACE2 of the WT, BA.1, and BA.2 (PDB).

