## Supplementary material for "Structure adaptation in Omicron SARS-CoV-2/hACE2: Biophysical origins of evolutionary driving forces": Key Measurements Made and Supplemental Table S1 and Figures S1-S7

Since its emergence, the COVID-19 pandemic has been sustained by a series of transmission waves initiated by new variants of the SARS-CoV-2 virus. Some of these arise with higher transmissivity and/or increased disease severity. Here we use molecular dynamics simulations to examine the modulation of the fundamental interactions between the receptor binding domain (RBD) of the spike glycoprotein and the host cell receptor (human angiotensin-converting enzyme 2: hACE2) arising from Omicron variant mutations (BA.1 and BA.2) relative to the original wild type strain. We find significant structural differences in the complexes which overall bring the spike protein and its receptor into closer proximity. These are consistent with and attributed to the higher positive charge on the RBD conferred by BA.1 and BA.2 mutations relative to the wild type. However, further differences between sub-variants BA.1 and BA.2 (which have equivalent RBD charges) are also evident: Mutations affect interdomain interactions between the up-chain and its clockwise neighbor chain, resulting in enhanced flexibility for BA.2. Consequently, additional close contacts arise in BA.2 which include binding to hACE2 by a second spike protein monomer, in addition to the up-chain - a motif not found in BA.1. Finally, the mechanism by which the glycans stabilize the up state of the spike protein differs for the wild type and the Omicrons. We also found the glycan on N90 of hACE2 turns from inhibiting, to facilitating the binding to Omicron spike protein. These structural and electrostatic differences offer further insight into the mechanisms by which viral mutations modulate host cell binding and provide a biophysical basis for evolutionary driving forces.

### Key Measurements Made

Trajectory data from the production run was obtained by taking snapshots of the system every 20 ps. As explained in the results the initial simulation structure evolves over the first part of the simulation. We therefore ignored the first 600 ns of the production run, and analysed the data from the last 400 ns of the simulation.

To study the interaction between amino acids we used the concept of close contact whereby a pair of residues was said to be in contact if the minimum distance between inter-residue atoms was less than 3 Å. We note that by doing so, we capture potential van-der-Waals, hydrogen bond and ionic attractions. The probability of occurrence of a close contact involving the residue was given as the number of frames in which the contact occurred divided by the total number of frames evaluated. Only events with probability higher than 50% are reported, unless otherwise noted.

To determine the chain domain $\alpha$'s compactness we calculated the radius of gyration, $R^{\alpha}_{gyr}$, using the non mass-weighted positions of every atom in the domain.

Similarly, the centre of mass (COM) of a chain domain is defined as the non mass-weighted average position using every atom of the domain. The distance between domains $\alpha$ and $\beta$, $d^{\text{ff}}_{\text{fi}}$, is then calculated as the length between their COM.

The RBD opening angle of chain A (as used by Fallon et al. (42)), $\theta_{\text{RBD-A}}$, is defined between the COMs of residues 338-517 of chain A, 324-327 and 538-585 of chain A, and 747-755 of chain C. By rotational symmetry, $\theta_{\text{RBD-C}}$ is defined the same as above but replacing A$\rightarrow$C and C$\rightarrow$B.

Root mean square fluctuation (RMSF) was calculated using VMD command *measure rmsf*, by comparing to the reference structure which is the time average over 600 ns to 1 $\mu$s

When studying glycans, the orientation of $\bar{\text{N}}90_{\text{hACE2}}$ with respect to S+hACE2 interface, was quantified by measuring the angle, $\Theta_{\text{N90}}$, made between the long axis defined by atom C4 on residue 6 with name BGAL, and atom C5 on residue 9 with name AFUC of $\bar{\text{N}}90_{\text{hACE2}}$ and the axis defined by residues 547 and 560 of hACE2 (see right panel of Figure S5).

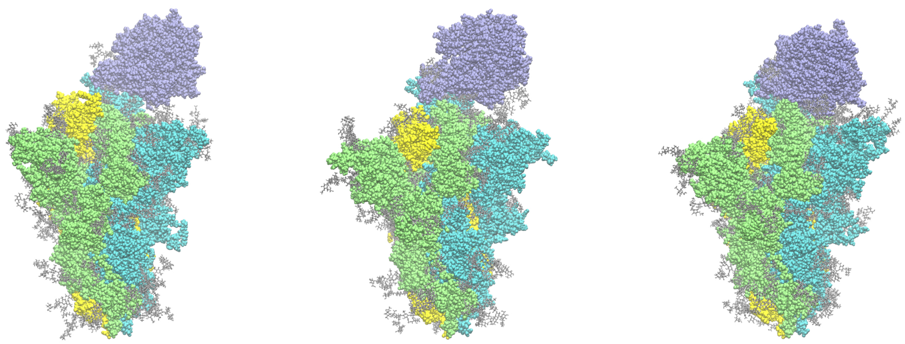

**Fig. S1.** Equilibrated structures of the S+hACE2. Left to right: WT, BA.1, BA.2; hACE2 in lilac, S-protein chain A in cyan, chain B in yellow, chain C in lime, and glycans in grey

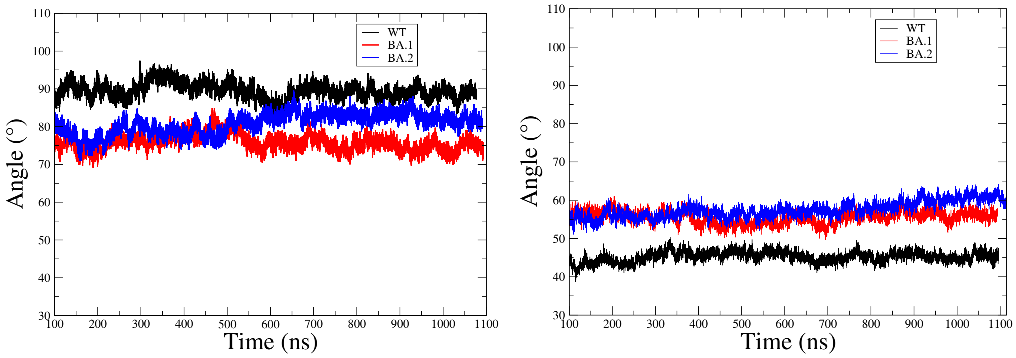


**Fig. S2.** Opening angle measurement of the up-chain (chain A: left) and its clockwise neighbor chain (chain C: right) of the S-protein

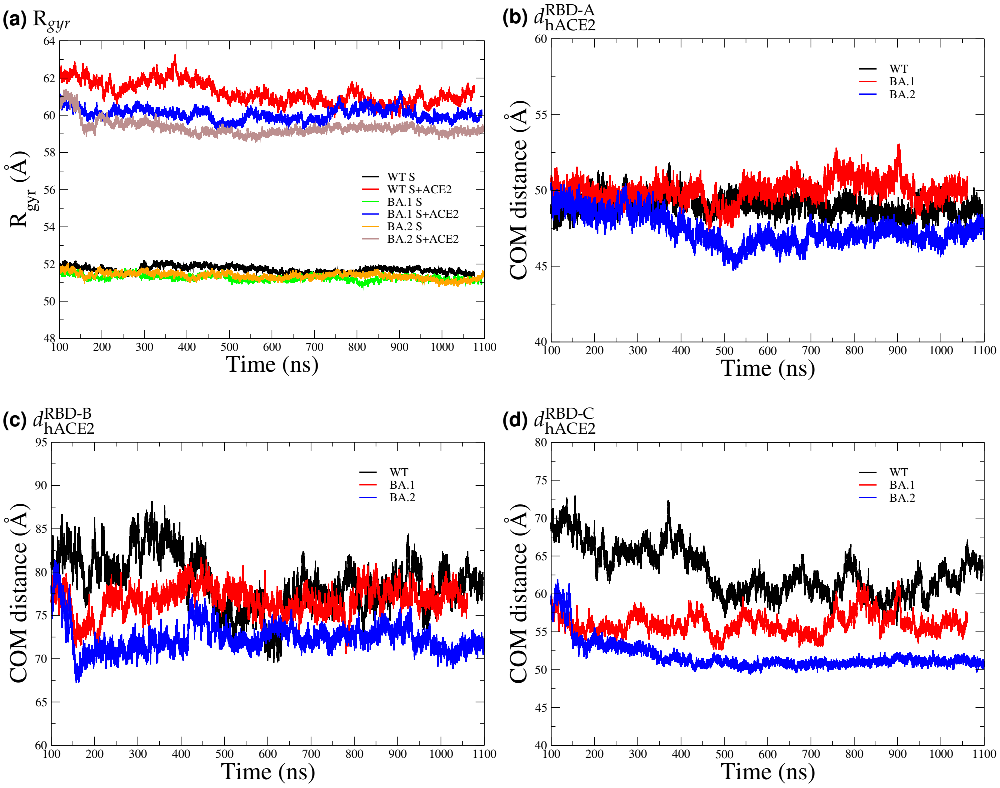


**Fig. S3.** (a) Radius of gyration of the up state trimeric S-protein; (b-d) Center of mass distances between hACE2 and the RBD of chains A, B, and C, respectively

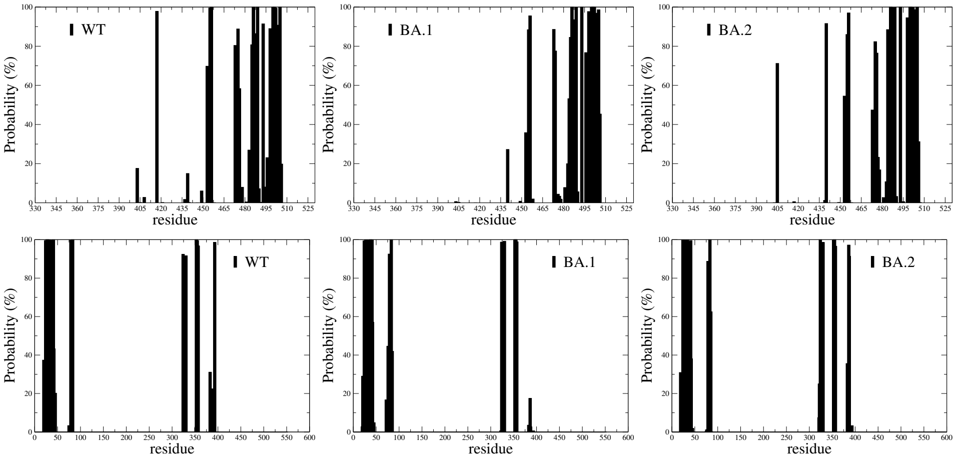


**Fig. S4.** Upper row: Probability of the residues on RBD-A come within 3 Å of hACE2; lower row: Probability of the residues on hACE2 come within 3 Å of RBD-A

**Table S1.** More key contact pairs (> 50% occurrence) RBD-A$\cdots$hACE2, including unmutated residues

| WT | BA.1 | BA.2 |
|---|---|---|
| F486$\cdots$Q24/L79/M82 | Q474$\cdots$Q24 | |
| | F486$\cdots$M85 | F486$\cdots$M85 |
| E484$\cdots$K31 | E484A$\cdots$Q76 | E484A$\cdots$L79 |
| G485$\cdots$L79 | G485$\cdots$L79 | |
| N501/Y505$\cdots$K353 | G496S$\cdots$K353 | N501Y/Q493R$\cdots$K353 |
| S477$\cdots$T20 | S477N$\cdots$Q24/T20 | S477N$\cdots$S19/T20 |
| T478$\cdots$Q24 | T478K$\cdots$Y83/P84 | T478K$\cdots$Q24/P84 |
| | | N440K$\cdots$E329 |
| | | D405N$\cdots$A387 |
| Q49$\cdots$E35/K31 | Q493R$\cdots$D38 | Q493R$\cdots$D38 |
| | | N501Y/Y505H$\cdots$K353 |
| Y505$\cdots$E37/R393 | | |

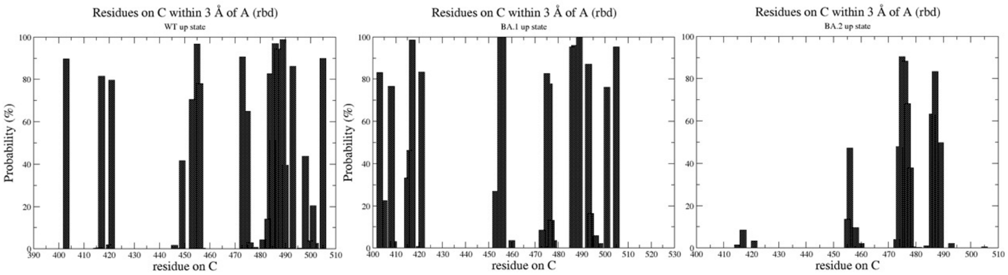


**Fig. S5.** Probability of close contacts by residues on RBD-C to RBD-A: clear reduction from WT to BA.1, and further reduction for BA.2, leaving RBD-C most flexible

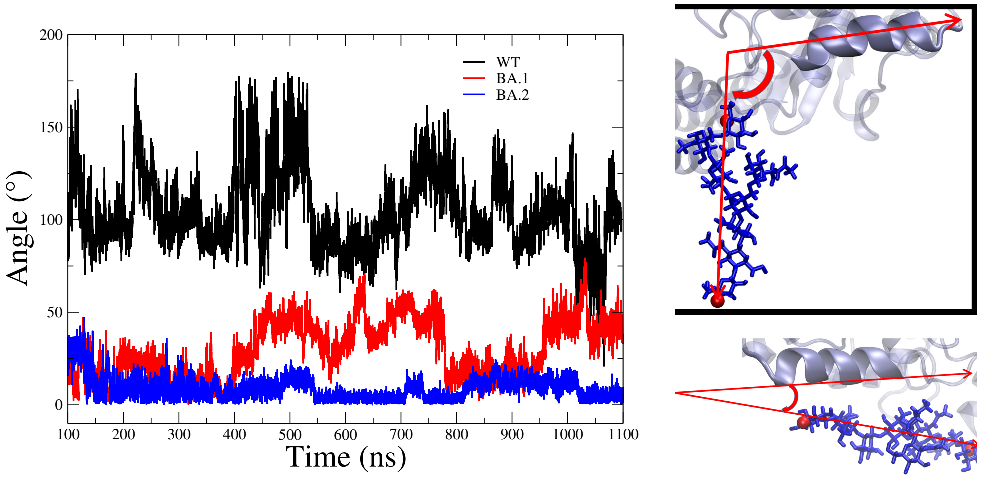


**Fig. S6.** Angle between the helix of hACE2 at the interface to S-protein, and the long axis of the glycan on its N90. Long axis is defined by atom C4 on residue 6 with name BGAL, and atom C5 on residue 9 with name AFUC

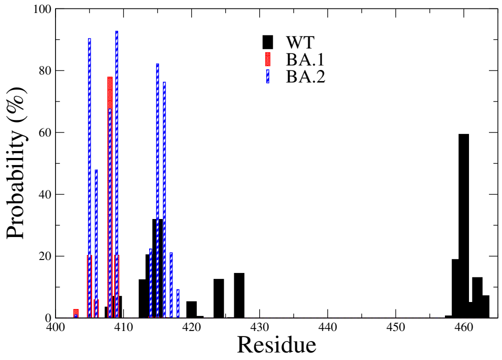


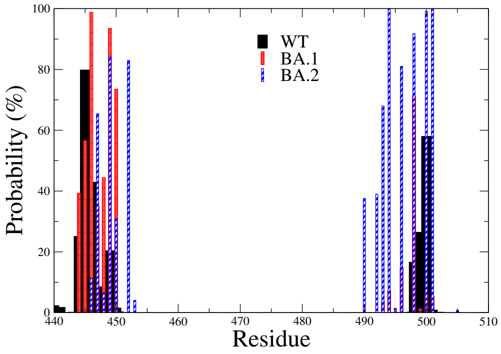


**Fig. S7.** Probability of (a) RBD-A (b) RBD-C residues in contact with glycan on N90 hACE2
